# Applying invasion criterion to cultural evolution

**DOI:** 10.1101/2025.11.12.688139

**Authors:** Shota Shibasaki, Masato Yamamichi

## Abstract

Cultural diversity is a fundamental aspect of human life and a driver of innovation; however, the underlying mechanisms remain poorly understood. While previous studies focused on how innovation generates cultural diversity, we investigated whether ecological theory, specifically the invasion criterion for understanding species coexistence in community ecology, could predict the maintenance of cultural diversity under various social learning biases. Using mathematical and individual-based models, we showed that the invasion criterion reliably predicts the duration of coexistence in the stochastic dynamics of multiple cultural traits under content-, conformity-, and anticonformity-biased social learning as well as under unbiased learning. The invasion criterion predicted moderate cultural coexistence under two types of model-biased social learning: prestige and success biases. In contrast, similarity-biased social learning, where people tend to learn from those with the same “tag” traits (e.g., gender, language), breaks the predictive power of the invasion criterion because the cultural traits were associated with the tag traits. Our results demonstrate that ecological theory offers valuable tools for understanding cultural evolution, but modifications are required to account for dynamic learning environments. Our model also showed that increasing the number of cultural models can prolong cultural coexistence under certain biases, with implications for fostering innovation and preserving culture. These findings bridge the ecological and cultural evolutionary theories and offer a unified framework for studying diversity maintenance across biological and cultural domains.

**Significance statement:** Cultural diversity benefits societal innovation and environmental adaptation, but how it is maintained remains unclear. This study applies the invasion criterion, an ecological concept, to predict how long different cultural traits coexist in a population. Simulations and mathematical analyses indicated that this ecological concept is appropriate for many types of social learning but fails for learning from similar individuals. Our mathematical analysis also showed that increasing the number of role models can support or hinder cultural diversity depending on the learning strategy. This work bridges biology and culture and offers new insights into how diversity persists.

## 1 Introduction

Culture is a “complex whole which includes knowledge, belief, art, morals, law, custom, and any other capabilities and habits” (Tylor, 1871/Tylor, 2010) and biological hallmark of humans. For about half a century, culture has been considered to follow Darwinian evolution, and cultural evolutionary theory has been developed to elucidate the complex dynamics of human culture (Boyd and Richerson, 2024; Fogarty et al., 2024). Various theories of cultural evolution have been adopted and extended from evolutionary biology (Claidière et al., 2014; Smolla et al., 2021). Diversity is a crucial topic in the evolution of human culture, as reflected in the more than 7,000 languages spoken worldwide (Eberhard et al., 2024; Bromham et al., 2024), and the increasing number of religions (Warf and Vincent, 2007; Lin et al., 2022).

Cultural diversity has mainly been investigated by comparing populations, i.e., cultural macroevolution (Mace and Jordan, 2011; Turchin et al., 2013). The macroevolutionary dynamics of human culture have been demonstrated in hunting tools (Peng and Nobayashi, 2021), language (Gray and Atkinson, 2003), and music (Savage, 2019). Although empirical studies showed that cultural diversity exists within populations (Nettle, 1998; Warf and Vincent, 2007), few have considered how it is maintained. Because variation is a driver of evolution, cultural evolutionary theories require mechanistic explanations for the maintenance of intrapopulation cultural variation.

This theoretical gap may partially stem from cultural evolution models traditionally relying on population genetics frameworks, which often assume “passive” or transient variation through mutation-selection balance (Loewe and Hill, 2010), although some population genetic studies focus on balancing selection (Ruzicka et al., 2025). For example, studies on the number of independent cultures (Strimling et al., 2009; Fogarty et al., 2015) assume increased cultural traits through innovation. In contrast, ecological theories consider the coexistence mechanisms that “actively” maintain species diversity despite resource competition (May, 1972; Chesson, 2000) without speciation (note that mutation is analogous to speciation in an ecological context and innovation in a cultural context). According to the invasion criterion in modern coexistence theory, the stable coexistence of diverse species requires that all in a community increase in abundance when each species is rare (Grainger et al., 2019). Many empirical ecological studies employ this theoretical framework (Letten et al., 2018; Shinohara et al., 2024).

Here, we applied the invasion criterion in cultural evolution to predict the coexistence of multiple cultures (instead of species) without innovation. Based on Schreiber et al. (2023), we expected the invasion criterion to reflect the duration of coexistence in a finite-sized population (instead of a community). This study focused on predicting the coexistence of multiple cultures that share roles (e.g., language, religion, or hunting tools) under unbiased and six major biased forms of social learning (Henrich and McElreath, 2003; Kendal et al., 2018): content, conformity, anticonformity, prestige, similarity, and success bias (Table 1). Content bias refers to the preference for cultural traits based on their inherent properties (Stubbersfield, 2022); e.g., some cultures are more likely to be inherited because they relate to social interaction (Mesoudi et al., 2006) or are minimally counterintuitive (Norenzayan et al., 2006). Conformity and anticonformity biases refer to frequency-dependent biases; thus, the contents of cultural traits do not affect the invasion criteria. The conformity bias makes major cultures more likely to be copied by others than their actual fractions, while making minor cultures less likely to be copied (Boyd and Richerson, 1985). Anticonformity biases reverse this relationship, making rare cultures more likely to be copied (Acerbi and Bentley, 2014). Under prestige, similarity, or success biases, learners inherit cultural traits based on models’ characteristics (i.e., model-biased social learning). Prestige bias occurs when individuals are more likely to learn from those regarded as more successful or knowledgeable (Henrich and Gil-White, 2001), even when the models cultural traits are unrelated to their success (Henrich and Broesch, 2011). Success bias allows learners to inherit cultural traits of those who have high fitness or payoffs (Baldini, 2012; Barrett et al., 2017). In this sense, success and prestige biases are connected because successful individuals are more likely to have high status. In this study, we used success bias when an individual’s survival was correlated with their cultural traits, whereas they were independent under prestige bias. Similarity bias indicates that individuals are more likely to learn from others who have similar characteristics to the learners, such as gender and spoken language (Buttelmann et al., 2013; Schniter et al., 2023). We refer to these traits as “tags.”

**Table 1:** Classification of social learning biases.

| Category | Subtype | Bias toward | Analytical solution | Predictability by invasion fitness |
| --- | --- | --- | --- | --- |
| 1. Unbiased | – | – | Yes | Yes |
| 2. Content bias | – | Attractive culture. | Yes | Yes |
| 3. Frequency-dependent bias | 3a. Conformity bias | Majority’s culture. | Yes | Yes |
|  | 3b. Anticonformity bias | Minority’s culture. | Yes | Yes |
| 4. Model bias | 4a. Prestige bias | Culture of prestigious models. | No | Partially |
|  | 4b. Success bias | Culture of successful models. | No | Partially |
|  | 4c. Similarity bias | Culture of similar models. | No | No |

Through the analysis of Markov chain models and individual-based simulations of cultural evolution in a finite population, we investigated whether the invasion criterion predicted the duration of the coexistence of multiple cultures under the social learning biases. This simple proxy can be helpful to estimate how the strength of social learning biases and the number of cultural models affect cultural diversity, which may facilitate societal innovation and animal conservation. We suggest that ecological theories provide valuable frameworks for elucidating cultural evolution; however, modifications and extensions are necessary because the invasion criterion reduced the predictability in the mode-based social learning.

## 2 Methods

### 2.1 Model description and calculation of invasion fitness

We assumed a population of fixed size *N* on a discrete-time scale, and each individual has one of multiple cultural traits. In other words, these cultures compete for individuals (Kandler and Laland, 2009). We focused on the cases of two cultural traits, *A* and *a*, and assumed that the two neutral cultural traits do not affect the mortality of individuals, except for the cases under success bias. In the Supporting Information, we relaxed these assumptions and analyzed three-culture and non-neutral two-culture models. The dynamics in this study obey the Moran process for calculation simplicity. At each time step, an individual is born, *K* individuals (1 ≤ *K* ≤ *N*) are randomly sampled as cultural models from *N* individuals, and a cultural trait is inherited from *K*. Subsequently, a randomly chosen individual from *N* individuals is replaced by a newborn individual. Suppose *N*_*A*_(*t*) is the number of individuals with culture *A* at time step *t*. Under the Moran process, the conditional probability of the number of individuals with culture *A* at time step *t* + 1, *P* (*N*_*A*_(*t* + 1) | *N*_*A*_(*t*)), depends on the mortality of an individual with *A* and the probability that the newborn individual inherits culture *A* through social learning *p*_*A*_(*N*_*A*_; *K*), which depends on *K* cultural models and social learning biases. Because the invasion criterion focuses on population growth when it is rare (Chesson, 2000), we focused on the case of *N*_*A*_(*t*) = 1. Although the ecological literature typically uses the term “invasion growth rate,” we use “invasion fitness” for consistency with evolutionary biology. Schreiber et al. (2023) showed that minimum invasion fitness among species predicts the coexistence time when the sum of total population size is finite. They defined invasion fitness as the expected population size of the focal rare species at the next time step.

Following their definition, we defined the invasion fitness of culture *A* as follows:

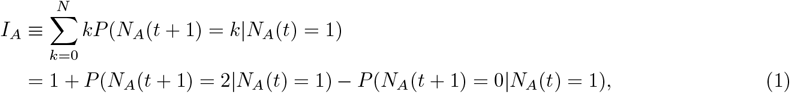

reflecting the Moran-process assumption (i.e., *N*_*A*_(*t* + 1) = {*N*_*A*_(*t*), *N*_*A*_(*t*) *±* 1*}*).

When *N*_*A*_(*t*) = 1, a newborn individual can learn culture *A* only if the unique individual with culture *A* is a member of *K* cultural models. This unique individual with culture *A* can be selected as a model with probability *K/N*. Then, the probability that a newborn individual inherits the rare culture *A* is expressed as follows:

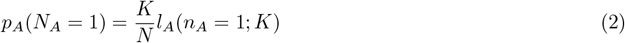

where *l*_*A*_(*n*_*A*_; *K*) represents the probability that a newborn individual learns culture *A* when the *n*_*A*_ cultural model possesses culture *A*. Social learning bias alters the formulation of *l*_*A*_(*n*_*A*_; *K*). The invasion fitness of culture *A* is simplified as follows:

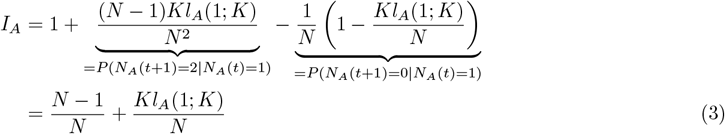

The invasion fitness of the other culture *I*_*a*_ is defined in a similar manner. For each scenario, we calculated the invasion fitness of the two cultural traits and used their minimum invasion fitness (MIF), min {*I*_*A*_*I*_*a*_}, as the potential indicator of the duration of coexistence.

### 2.2 Formulation of social learning biases

#### 2.2.1 Absence of social learning biases

In the baseline model, an individual randomly copies a cultural trait from the cultural models, representing an unbiased social learning scenario. The probability that a newborn individual learns culture *A* depends solely on the number of individuals with culture *A* within *K* cultural models.

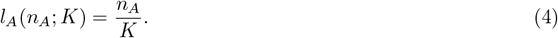

By substituting the above *l*_*A*_(*n*_*A*_; *K*) for Eq (3), we find that MIF in the absence of social learning biases is always one. Therefore, we used this scenario as a baseline for the duration of cultural coexistence in the below scenarios.

#### 2.2.2 Content bias

Content bias indicates that two cultures differ in cultural preferences. Without loss of generality, we defined the preference for culture *a* as one, and that for culture *A* as 1 + *s* (*s* ≥ 0), where *s* represents the strength of the content bias. Then, *l*_*A*_(*n*_*A*_; *K*) is expressed as follows:

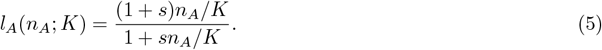

It is obvious that *s* = 0 returns to the unbiased social learning in Eq (4). We focused on the cases of *s* = 0.1, 0.5, 1, 1.5, 2. In the presence of three or more cultural traits, the denominator of Eq (5) becomes ∑_*i*_(1 + *s*_*i*_)*n*_*i*_*/K* where *s*_*i*_ represents the bias toward culture *i* and *n*_*i*_ is the number of cultural models with culture *i*.

#### 2.2.3 Conformity and anticonformity biases

The conformity and anticonformity biases represent positive and negative frequency-dependent learning biases, respectively. Under (anti)conformity bias, an individual is more (less) likely to learn about a majority culture than its fraction within *K* cultural models (Boyd and Richerson, 1985). We implemented the (anti)conformity bias following Nöbel et al. (2023).

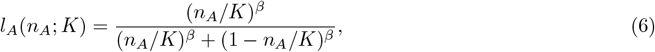

where *β* is a real parameter that determines whether this function represents conformity or anticonformity bias and its strength.

Eq (6) has the following advantages: First, multiple learning biases can be represented by tuning a single parameter *β*; *β >* 1 represents the conformity bias, *β* = 1 returns to unbiased social learning, as in Eq (4), and *β <* 0 corresponds to the anticonformity bias. When 0 *< β <* 1, the function represents a “weak-conformist” bias (Claidière et al., 2012); however, we did not consider this bias because we followed the definition of conformist bias in Boyd and Richerson (1985). Second, Eq (6) can easily be extended to include more than two cultural traits by rewriting the denominator as ∑_*i*_(*n*_*i*_*/K*)^*ω*^.

We slightly modified Eq (6) when implementing the anticonformity bias. Because we did not consider mutations in this study, an individual cannot learn culture *A* (*a*) when none of the models have that culture. Thus, the probability that an individual learns culture *A* under the anticonformity bias is zero (one) when none (all) of his/her cultural models have trait *A*. We used *β* = 1.1, 1.5, 2, 2.5, 3 for the conformity bias and *β* = −1.1, −1.5, −2, *−*2.5, *−*3 for the anticonformity bias.

#### 2.2.4 Prestige bias

Prestige bias indicates the likelihood of culture being copied from those with higher prestige. In other words, the probability that a newborn individual learns depends on the prestige of the models with culture *A*. Because this study investigated whether the invasion criterion predicts cultural coexistence, we focused on the minimal binary prestige model (Acerbi et al., 2022). We assumed that each individual had either low or high prestige and that a newborn individual was more likely to learn from high-prestige cultural models than from low-prestige ones. Without loss of generality, we set the learning bias toward the low-prestige cultural models as one and that toward the high-prestige cultural models as 1 + *W* (*W* 0), where *W* represents the strength of the prestige bias (*W* = 0, 0.1, 1, 10, 100). We assumed that a newborn individual inherited the prestige of the dead individual at each time step so that the fractions of low- and high-prestige individuals were constant over time. We analyzed cases in which the fractions of high-prestige individuals were 0.1, 0.2, and 0.5.

Invasion fitness depends on the probability that either a low-prestige or high-prestige individual holds a focal rare culture. In SI 1, we calculated the conditional probability that a newborn individual learns the focal rare culture from low- and high-prestige individuals. From this, we derived the conditional invasion fitness when rare culture *A* is held by a low-prestige individual:

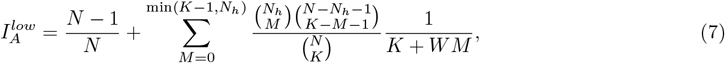

where *N*_*h*_ is the number of high-prestige individuals in the population, and *M* is the number of high-prestige individuals within *K* cultural models. From this conditional invasion fitness, we can obtain the conditional MIF given that a low-prestige individual holds a focal rare culture.

We confirmed that the invasion fitness was always one when naïvely assuming that the probability of becoming a holder of the rare culture was 1*/N* for all individuals. See SI 1 for mathematical details.

#### 2.2.5 Success bias

Under success bias, newborn individuals are more likely to learn cultural traits from those with high fitness or payo!. Because the population size *N* was fixed in our model, we assumed that a newborn individual with success bias was more likely to learn from cultural models with lower mortality rates. To implement the cultural impact on the mortality rate, we allowed cultural trait *A* to increase or decrease the mortality of its owner while fixing the mortality rate of individuals with cultural trait *a* as one, which returns to a special case of content-biased social learning in which individuals prefer culture *A* to *a* and culture *A* decreases the mortality rate of its owner (see SI 2).

To highlight the difference between content and success biases, we analyzed the success bias under environmental fluctuations to temporarily change the success of cultural traits. Suppose that the two cultural traits represent different hunting tools, and that each tool is suitable for hunting different prey, the degree of success of each culture depends on the relative abundance of the prey. Considering this, we analyzed the minimal model of success-biased social learning in a fluctuating environment. For simplicity, we assumed that the environment is represented by a dichotomous random variable (*ξ*(*t*) = ±1), mortality rate of (and preference for) individuals with cultural trait *A* depends on the environmental condition, increase in mortality rate by *W* results in a decrease in the preference for culture *A* by *W* and vice versa (*W* = 0.1, 0.25, 0.5, 0.75, 0.9), and mortality rate of (and preference for) individuals with cultural trait *a* is independent of the environmental conditions. The environmental condition changed from *ξ* to *ξ* at *τ* step, where τ follows a geometric distribution with a mean of 1*/σ* and σ = 1, 5, 10, 50, 100 represents the environmental stability.

The invasion fitness of the two cultural traits in environment *ξ* is expressed as follows:

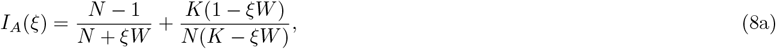

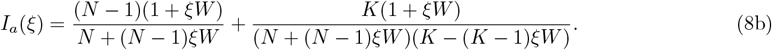

Here, *W* = 0 returns to unbiased social learning because the invasion fitness becomes one. The expected invasion fitness of each cultural trait was given as the average invasion fitness weighted by the stationary distribution of *ξ*. Because we assumed a symmetric environmental switch, the stationary distribution is *P* (*ξ* = 1) = *P* (*ξ* = −1) = 0.5. Thus, the MIF is the minimum value of the expected invasion fitness for the two cultural traits.

#### 2.2.6 Similarity bias

Under similarity bias, the tag traits (e.g., gender) of a newborn and cultural model affect the probability that the newborn learns from the models. Without loss of generality, we assumed that a newborn individual learned from a cultural model with an identical or unmatched tag trait at a rate of 1 + *W* and one, respectively. Here, *W* represents the strength of the similarity bias (*W* = 0, 0.1, 1, 10, 100).

We assumed that the population had binary tag traits, 0 and 1, and that the number of individuals with these markers was even. This situation corresponds to, for example, gender-biased learning under the unbiased sex ratio. We used *N*_0_ to denote the number of individuals with a tag of 0. We fixed the fractions of the two tag traits to 0.5 so that neither tag would become extinct owing to random drift through the simulations. Accordingly, a newborn individual at time *t* inherits the tag trait of a dead individual at that time.

When only a single individual has culture *A* (i.e., *N*_*A*_(*t*) = 1), and this individual has tag trait of 0, the probability that a newborn individual learns culture *A* is as follows:

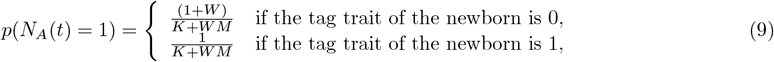

where *M* ≥ 1 represents the number of individuals with tag trait 0 within *K* cultural models. Because the number of individuals with cultural trait *A* increases from one only if a single individual does not die and a newborn individual (either with tag trait 0 or 1) learns the cultural trait, the probability is expressed as follows:

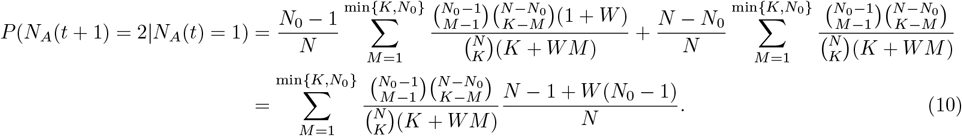

Similarly, cultural trait *A* becomes extinct if the owner dies and no newborn (with tag trait 0) learns it.

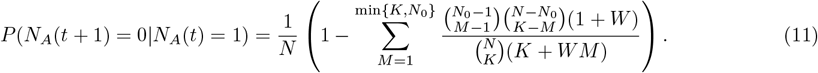

When the rare culture *A* is owned by an individual with tag trait 1, we can perform a similar calculation by replacing *N*_0_ with *N* − *N*_0_. Because we assumed *N*_0_ = *N/*2, and individuals of both tag traits are equally likely to be owners of the rare culture *A*, the invasion fitness of culture *A* under such a similarity bias is expressed as follows:

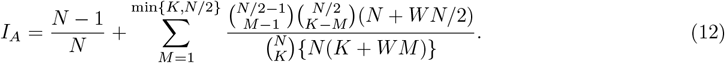

Similarity bias with *W* = 0 returns to the unbiased social learning because *I*_*A*_ = 1.

### 2.3 Calculating the duration of coexistence

We investigated the mean duration of coexistence (i.e., time until one cultural trait goes extinct) using either Markov chain or individual-based models. In the two-culture trait scenarios, the mean duration of coexistence is identical to the mean time until absorption in any absorbing state (Otto and Day, 2007). Here, we refer to “mean duration of coexistence” as it is different from the mean time until absorption when three or more cultural traits exist in a population: The duration of coexistence measures the time until one cultural trait goes extinct, while the time until absorption measures the time until only one cultural trait remains in a population. Under unbiased, content-biased, and (anti)conformity-biased social learning, we used the Markov chain model, because *P* (*N*_*A*_(*t* + 1)|*N*_*A*_(*t*)) depends on the number of individuals with culture *A*. In the remaining scenarios, we used the individual-based model because *P* (*N*_*A*_(*t* + 1) |*N*_*A*_(*t*)) depends on who has culture *A*, which complicates calculating the transition matrix.

In the Markov chain model, the transition matrix *P* is defined as follows:

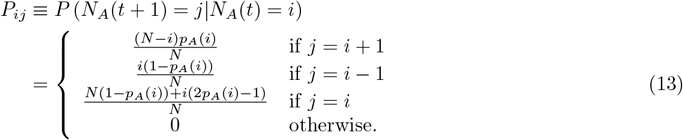

In SI 2, we considered the case in which culture *A* affects owner mortality rate.

In the Markov chain model, we derived the mean duration of cultural coexistence based on the initial probability distribution of the number of individuals in culture *A P* (*N*_*A*_(0)) (Otto and Day, 2007). We assumed that each individual initially had culture *A* with a probability of 0.5 (otherwise, they had culture *a*). Thus, *P* (*N*_*A*_(0)) follows a binomial distribution with mean *N/*2. We obtained the mean duration of coexistence under unbiased, content-biased, and (anti)conformity-biased social learning when *N* = 10, 20, 30 and *K* = 3, 5, 7.

In the individual-based model, we analyzed cases of prestige-, success-, and similarity-biased social learning. As in the above cases, each individual initially has culture *A* with a probability of 0.5 (otherwise, they have culture *a*). For each of the three biases, the strength *W*, population size *N*, and number of cultural models *K* (and fraction of high-prestige individuals under the prestige bias or environmental stability *σ* under the success bias), we ran 1,000 replicates, and derived the mean duration of coexistence and its standard error.

### 2.4 Applying the invasion criterion to cultural evolution

Because the invasion fitness is always one under the unbiased social learning (see Eq (4)), we used the mean duration of coexistence of under the unbiased social learning as the baseline. We examined the *qualitative* and *quantitative* prediction of the invasion criterion as follows:

#### Qualitative prediction

If the MIF among the cultural traits is lower (higher) than one, the mean duration of cultural coexistence is shorter (longer) than the cases under the unbiased social learning.

#### Quantitative prediction

The MIF among the cultural traits positively correlates with the mean duration of cultural coexistence.

The invasion criterion qualitatively predicts the coexistence time in ecological scenarios but often fails in quantitative predictions (Pande et al., 2020; Ellner et al., 2020). We performed regression analysis to assess quantitative predictive performance. For the anticonformity bias, we fit the log_10_ mean duration of coexistence to the linear model because it seemed to increase exponentially over the MIF. For the other five social learning biases, the following logistic curve was used:

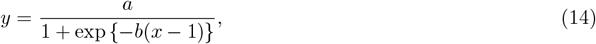

where *x* is the MIF, *y* is the mean duration of coexistence, and *a* and *b* are logistic curve parameters. Note that *y* = *a/*2 when *x* = 1: i.e., the mean duration of coexistence is half that of *a* when the MIF is one (under unbiased social learning). We used a logistic curve because the five biases return to unbiased social learning when their strength is zero and the mean duration of coexistence is expected to converge to that of unbiased social learning. We used Scipy version 1.15.3 (Virtanen et al., 2020) to estimate the parameter values; specifically, the linregress function for the linear model and curve fit function for the logistic regression. We evaluated the goodness of fit of the statistical models based on the (pseudo-)*R*^2^ and MAPE. When we fitted the data under content- and conformity-biased social learning, we also included the data on unbiased social learning because it corresponds to cases of minimum bias strength.

## 3 Results

### 3.1 Social learning biases alter invasion fitness

We first visualized how the minimum invasion fitness (MIF) of the two cultural traits changed with the strength of the social learning biases, population size *N*, and number of cultural models *K* (see 2.2 for mathematical details). MIF monotonically decreased with the strength of content-biased social learning (Fig. 1A) because the more substantial bias increases the invasion fitness of one culture while decreasing the other. The strength of the conformity and anticonformity biases decreased and increased the MIF monotonically, respectively (Fig. 1B and C), indicating that the (anti)conformity bias decreases (increases) the probability that a newborn inherits a rarer cultural trait. The strength of the prestige bias monotonically decreased the conditional MIF when the focal rare cultural trait was held by a low-prestige individual because a newborn is more likely to learn from high-prestige people under greater prestige bias (Fig. 1D). Under success bias, we assumed that a more substantial bias resulted in larger fluctuations in the preference for one culture and the mortality of its owners. Although the environmental conditions altered the invasion fitness of the two cultural traits, the success bias strength monotonically decreased the MIF (Fig. 1F). The similarity bias strength first decreased the MIF but then increased it upon exceeding a certain threshold.

**Figure 1:**
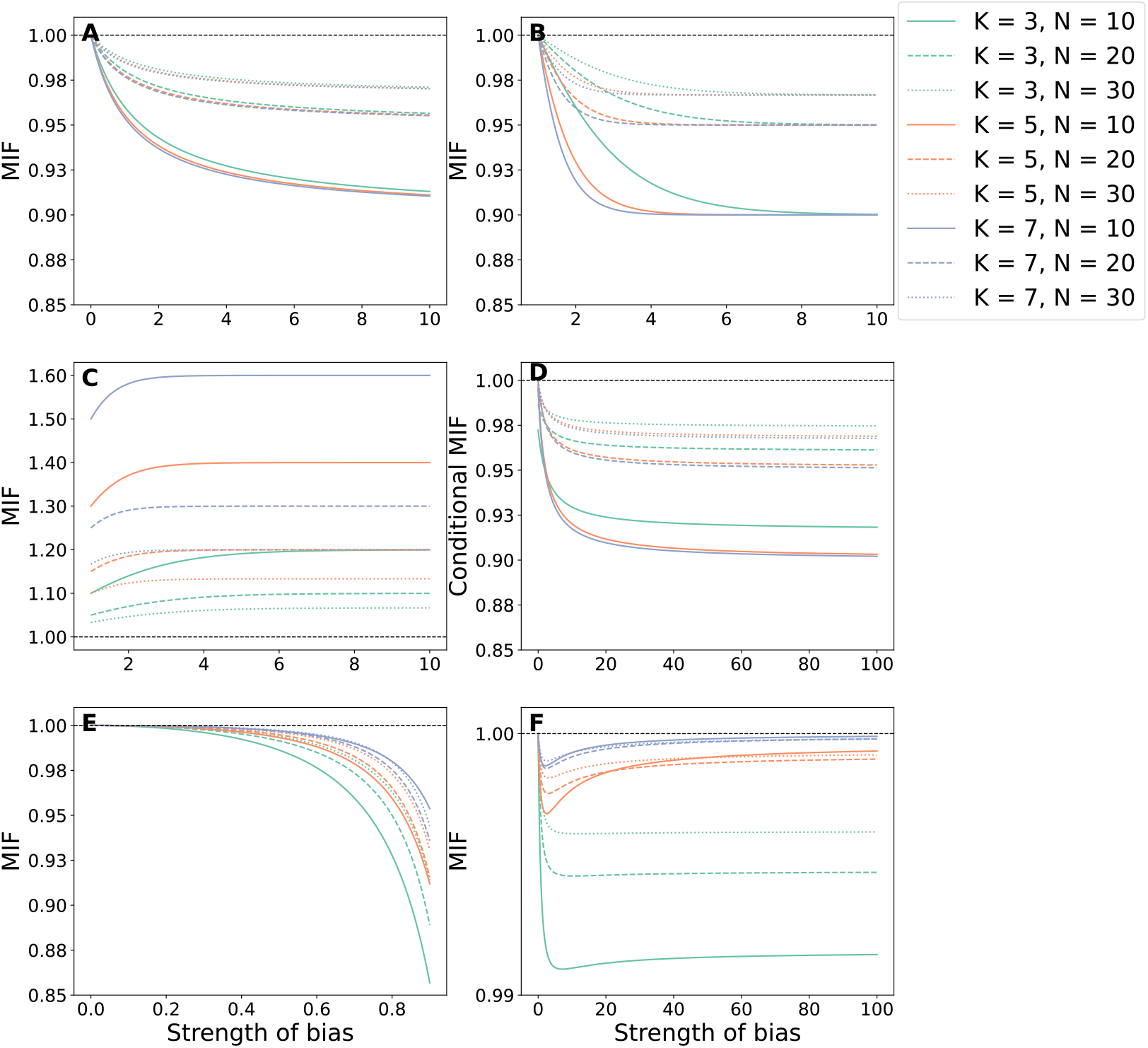
Social learning biases change the minimum invasion fitness The minimum invasion fitness (MIF) under the six social learning biases are shown (A: content, B: conformity-, C: anticonformity-, D: prestige-, E: success-, and F: similarity-biased social learning). The colors of the lines represent the number of cultural models *K*, and the line styles represent the population size *N*. The black dashed lines represent the MIF of one, which corresponds to the unbiased social learning scenario. Panel D sets the number of high-prestige people as *N*_*h*_ = *N/*2.

Fig. 1 shows that the population size *N* and number of cultural models *K* altered the MIF. Under all the social learning biases, the MIF was closer to one with increasing population sizes. This corresponds to the first term in Eq (3), (*N −* 1)*/N*, and a similar term for success-biased social learning (see 2.2). However, the impact of the number of cultural models on MIF differed among the types of biases. A larger number of cultural models decreased MIF under content-, conformity-, and similarity-biased social learning. In contrast, the number of cultural models increased the MIF under anticonformity-, success-, and similarity-biased social learning. These results indicate that the MIF summarizes the details of social learning.

### 3.2 Invasion criterion predicts the cultural coexistence under no social learning bias, content bias, and frequency-dependent biases

We investigated the duration for which the two competing neutral cultures *A* and *a* coexisted under unbiased, content-biased, and frequency-dependent (conformity and anticonformity) social learning. These scenarios serve to determine the applicability of the invasion criterion for predicting the cultural coexistence duration under the Markov chain model framework (Otto and Day, 2007). Under unbiased social learning (Fig. 2), the MIF is one regardless of the number of cultural models *K* and population size *N*. However, the duration of coexistence under the Moran process generally depends on population size. Therefore, we set unbiased scenarios as baselines for each population size (Figs 2A and D: *N* = 10, Figs 2B and E: *N* = 20, and Figs 2C and F: *N* = 30). Under content or conformity bias, the MIF was always lower than one, and the two cultural traits coexisted for a shorter duration than in the unbiased cases (Figs.2D–F). The anticonformity bias increased the MIF, and the coexistence time increased exponentially with increasing MIF (Figs. 2A–C).

**Figure 2:**
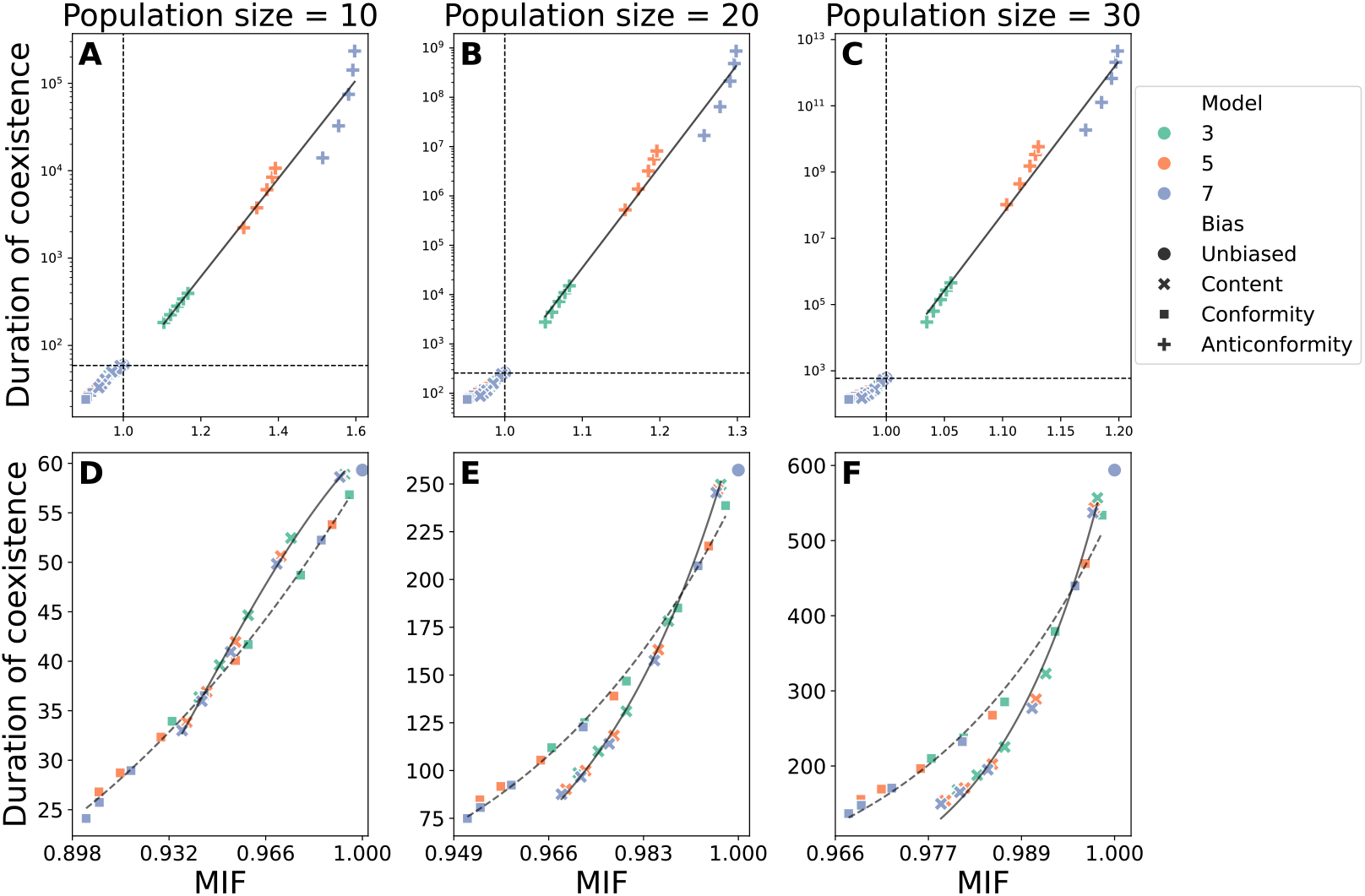
Minimum invasion fitness predicts the cultural coexistence under unbiased, content-biased, and frequency-dependent biased social learning

Under these three biases, the MIF precisely predicted the mean duration of coexistence. Table 2 shows that all statistical models had mean absolute percentage error (MAPE) lower than approximately 10%. The statistical model revealed a difference between content- and conformity-biased social learning: Although they produce similar MIF values and coexistence durations, they differ in how the MIF affects the duration. The duration of coexistence increased more steeply under content bias than under conformity bias; however, these biases returned to the unbiased scenario when the strength of the learning bias was zero and one, respectively. Consequently, the two cultural traits coexisted longer under content bias than under conformity bias when the MIF was close to one, whereas they coexisted longer under conformity bias than under content bias when the MIF was small.

**Table 2:** Summary of regression analysis.

| Bias | $N$ | Model | (pseudo-)$R^2$ | MAPE |
| --- | --- | --- | --- | --- |
| Content | 10 | $y = \frac{122.36}{1+\exp\{-14.81(x-1)\}}$ | 0.98 | 2.34 |
| | 20 | $y = \frac{526.06}{1+\exp\{-53.25(x-1)\}}$ | 0.99 | 3.05 |
| | 30 | $y = \frac{1196.34}{1+\exp\{-103.6(x-1)\}}$ | 0.97 | 8.78 |
| Conformity | 10 | $y = \frac{117}{1+\exp\{-13.77(x-1)\}}$ | 1.00 | 1.62 |
| | 20 | $y = \frac{495.44}{1+\exp\{-37.56(x-1)\}}$ | 0.98 | 5.49 |
| | 30 | $y = \frac{1121.77}{1+\exp\{68(x-1)\}}$ | 0.96 | 10.01 |
| Anticonformity | 10 | $\log_{10} y = 5.63x - 3.96$ | 0.97 | 2.51 |
| | 20 | $\log_{10} y = 20.6x - 18.12$ | 0.98 | 2.93 |
| | 30 | $\log_{10} y = 46.20x - 43.09$ | 0.98 | 3.93 |
| Prestige | 10 | $y = \frac{126.17}{1+\exp\{-9.33(x-1)\}}$ | 0.54 | 13.95 |
| | 20 | $y = \frac{522.39}{1+\exp\{-24.6(x-1)\}}$ | 0.73 | 12.36 |
| | 30 | $y = \frac{120.734}{1+\exp\{-39.48(x-1)\}}$ | 0.74 | 13.99 |
| Success | 10 | $y = \frac{97.47}{1+\exp\{-14.50(x-1)\}}$ | 0.39 | 36.32 |
| | 20 | $y = \frac{412.12}{1+\exp\{-24.71(x-1)\}}$ | 0.47 | 50.11 |
| | 30 | $y = \frac{985.69}{1+\exp\{-59.98(x-1)\}}$ | 0.53 | 48.72 |

$x$ : MIF, $y$ : mean duration of coexistence.

Overall, the Markov chain models indicated that the invasion criterion qualitatively predicts the duration of cultural coexistence under unbiased, content-biased, conformity-biased, and anticonformity-biased social learning. When fixing the social learning bias type, the statistical analysis confirmed that the invasion criterion quantitatively predicted the duration of coexistence. However, invasion criteria cannot precisely predict the duration of coexistence across social learning biases.

In the Supporting Information, we examined the robustness of the above results in two extended scenarios. First, we analyzed cases in which the mortality rates of individuals depended on cultural traits (i.e., the two cultural traits were not neutral). Such a model extension did not change the relationship between MIF and the duration of coexistence (Fig. S1 and Table S1). Furthermore, we extended our model to three cultural traits. Fig. S3 and Table S2 show that the MIF precisely predicted the coexistence of the three cultural traits under unbiased, content-biased, conformity-biased, and anticonformity-biased social learning (MAPE was at most 13.66%). These results highlight the robustness of the invasion criterion for predicting cultural coexistence under social learning biases.

### 3.3 Invasion fitness partially predicts the cultural coexistence under prestige and success biases but not under similarity bias

We investigated whether the MIF predicted cultural coexistence under prestige-, success-, and similarity-biased social learning. Under these model biases, the probability that a newborn learns either of the two cultural traits depends on the owner rather than the number of individuals. Because this complicates the Markov chain analysis, we used an individual-based model.

The invasion fitness and the mean duration of the coexistence of the two cultural traits under unbiased (circles), content bias (x marks), conformity bias (squares), and anticonformity bias (cross marks) social learning. The colors of the symbols represent the number of cultural models *K*, and each panel differs in the population size *N*. A: *N* = 10, B: *N* = 20, and C: *N* = 30. Panels D, E, and F correspond to the lower-left parts (i.e., the invasion fitness is smaller than one and the duration of coexistence is shorter than that under the unbiased social learning scenarios) of panels A, B, and C, respectively. The solid lines on panels A, B, and C show the regression lines to the anticonformity bias. The solid and dotted lines on panels D, E, and F represent the regression lines to the logistic curves of the content- and conformity-social learning biases, respectively.

The conditional invasion fitness (invasion fitness when the a low-prestige individual has a rare culture) and the mean duration of the coexistence of the two cultural trait under the binary prestige bias. The colors of the symbols represent the number of cultural models *K*, the error bars show the standard deviation, and each panel differs in the population size *N*. A: *N* = 10, B: *N* = 20, and C: *N* = 30. The horizontal dashed lines represent the mean duration of coexistence without social learning biases (Fig. 2). The black lines represents the fitted logistic curves.

#### 3.3.1 Prestige bias

We implemented a binary prestige bias, where each individual had either low or high prestige. High-prestige people are more likely to teach a newborn *W* (*>* 0) than low-prestige people, reflecting the strength of the prestige bias. To calculate MIF when using binary prestige bias, one needs to know how often low- and high-prestige individuals hold each cultural trait when only one individual holds the focal cultural trait. However, this probability cannot be obtained without simulations. Therefore, we calculated two types of invasion fitness as proxies.

Naïve invasion fitness assumes that each individual has an identical probability of being a holder of a rare cultural trait; in other words, a rare culture is held by low- and high-prestige people following their fractions in the population, respectively. In SI 1, we show that the naïve invasion fitness is always one under the prestige bias regardless of the fraction of prestige people, population size, number of cultural models, and bias strength. Because the mean coexistence time varied (see below), this proxy was not useful.

Conditional invasion fitness occurs when a low-prestige individual holds a focal rare culture. Fig. 3 shows that the conditional MIF was positively correlated with the mean duration of coexistence. However, the deviation from the logistic curve was large under prestige bias. The pseudo-*R*^2^ was smaller and MAPE was larger than under content-, conformity-, and anticonformity-biased social learning (Table 2). In other words, conditional MIF can quantitatively predict cultural coexistence under a simple prestige bias to some degree, but the accuracy is lower than that under content, conformity, and anticonformity biases. In some cases, the mean duration of coexistence slightly (less than 10%) exceeded that of the unbiased scenarios (dashed lines in Fig 3), while the conditional invasion fitness was lower than one. This indicates that the MIF failed to qualitatively predict the duration of coexistence.

**Figure 3:**
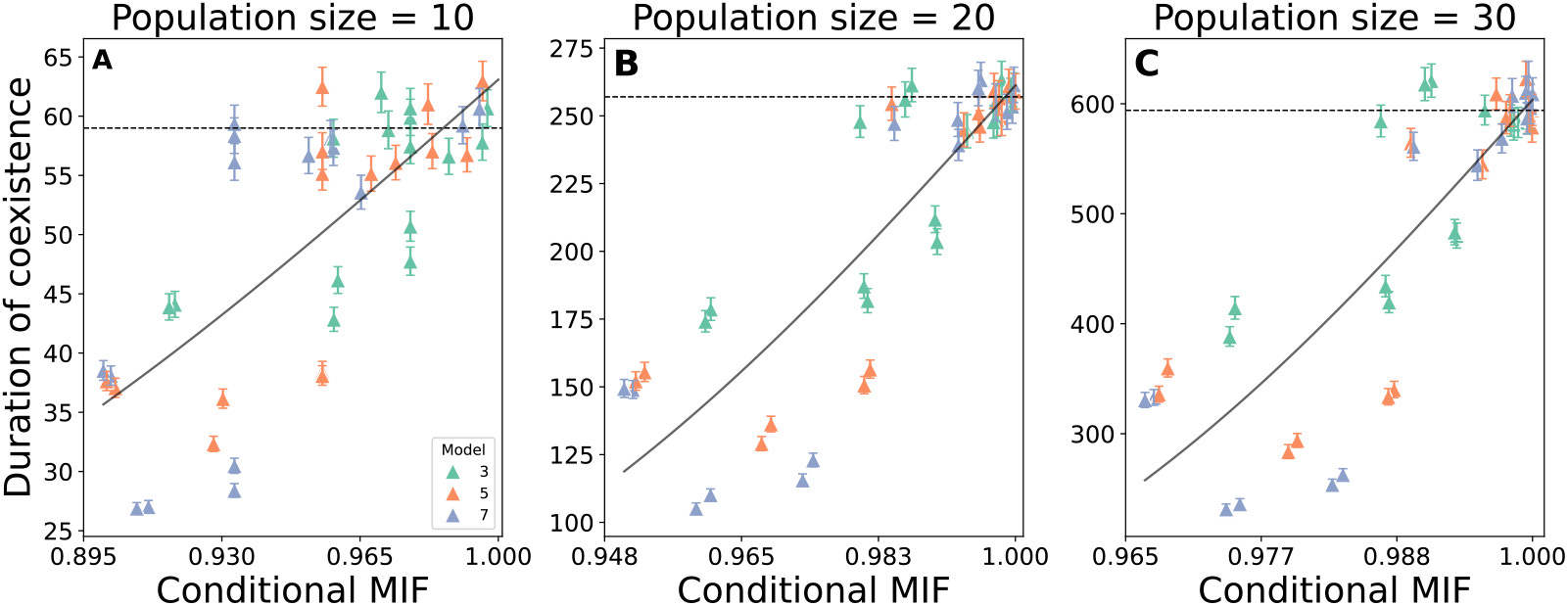
Conditional minimal invasion fitness moderately predicts the duration of coexistence under the prestige bias

#### 3.3.2 Success bias

Under success bias, a cultural trait is more likely to be learned from others if its holder has a higher fitness or payo!. Because our simulations followed the Moran process, the lower mortality rate implied higher fitness. We analyzed a scenario in which environmental conditions fluctuated over time. For simplicity, we assumed that environmental conditions affected the mortality of individuals with cultural trait *A* (and the social learning bias toward such people), whereas environmental fluctuations did not affect cultural trait *a*. Fig. 4 shows that the MIF was positively correlated with the mean duration of coexistence. In addition, both the MIF and mean duration of coexistence were smaller than those under unbiased simulations (dashed lines in Fig. 4). However, the MIF was independent of environmental stability (the inverse of the environmental fluctuation rate), whereas it altered the mean duration of coexistence (Fig. S3). This resulted in a small pseudo-*R*^2^ and large MAPE of the logistic curves in Table 2. However, fixing the environmental stability can improve pseudo-*R*^2^ and MAPE (Table S3). Overall, while the qualitative prediction by the invasion criterion was supported in success-biased social learning with environmental fluctuations, it cannot incorporate the change in the duration of coexistence over environmental stability.

**Figure 4:**
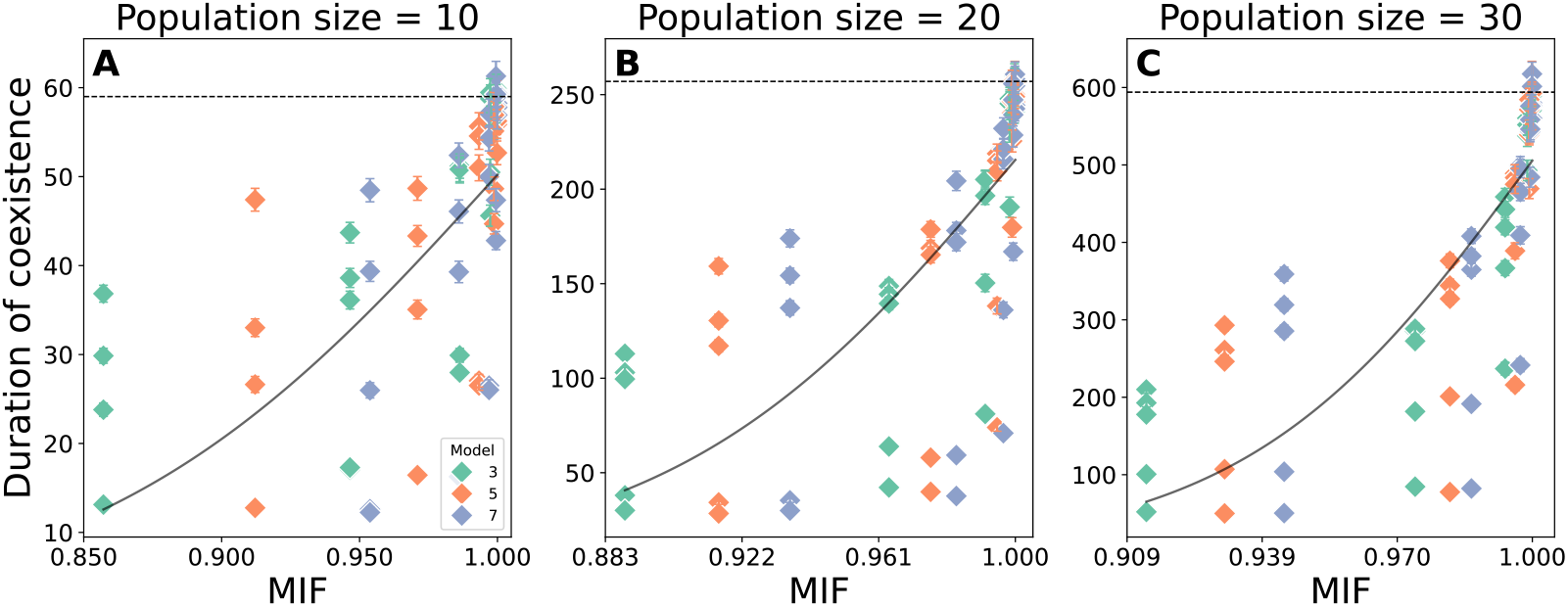
Invasion fitness moderately correlates with the duration of coexistence under the success bias The expected invasion fitness and the mean duration of the coexistence of the two cultural traits under the success bias with environmental fluctuations. The colors of the symbols represent the number of cultural models *K*, the error bars show the standard deviation (some of them are too small to be seen), and each panel differs in the population size *N*. A: *N* = 10, B: *N* = 20, and C: *N* = 30. The horizontal dashed lines represent the mean duration of coexistence without social learning biases (Fig. 2). The black solid lines represent the fitted logistic curves.

#### 3.3.3 Similarity bias

Under similarity bias, each individual has cultural traits and tags. In the individual-based simulations, we implemented a binary tag (0 and 1), whose fraction was even in the population, and assumed that a newborn individual was more likely to learn from models with the same tag and similarity bias strength *W*. However, Fig. 5A–C shows that the MIF was not correlated with the mean duration of coexistence under the similarity bias. In addition, while the invasion fitness was slightly lower than one, the two cultural traits coexisted longer than under unbiased social learning.

**Figure 5:**
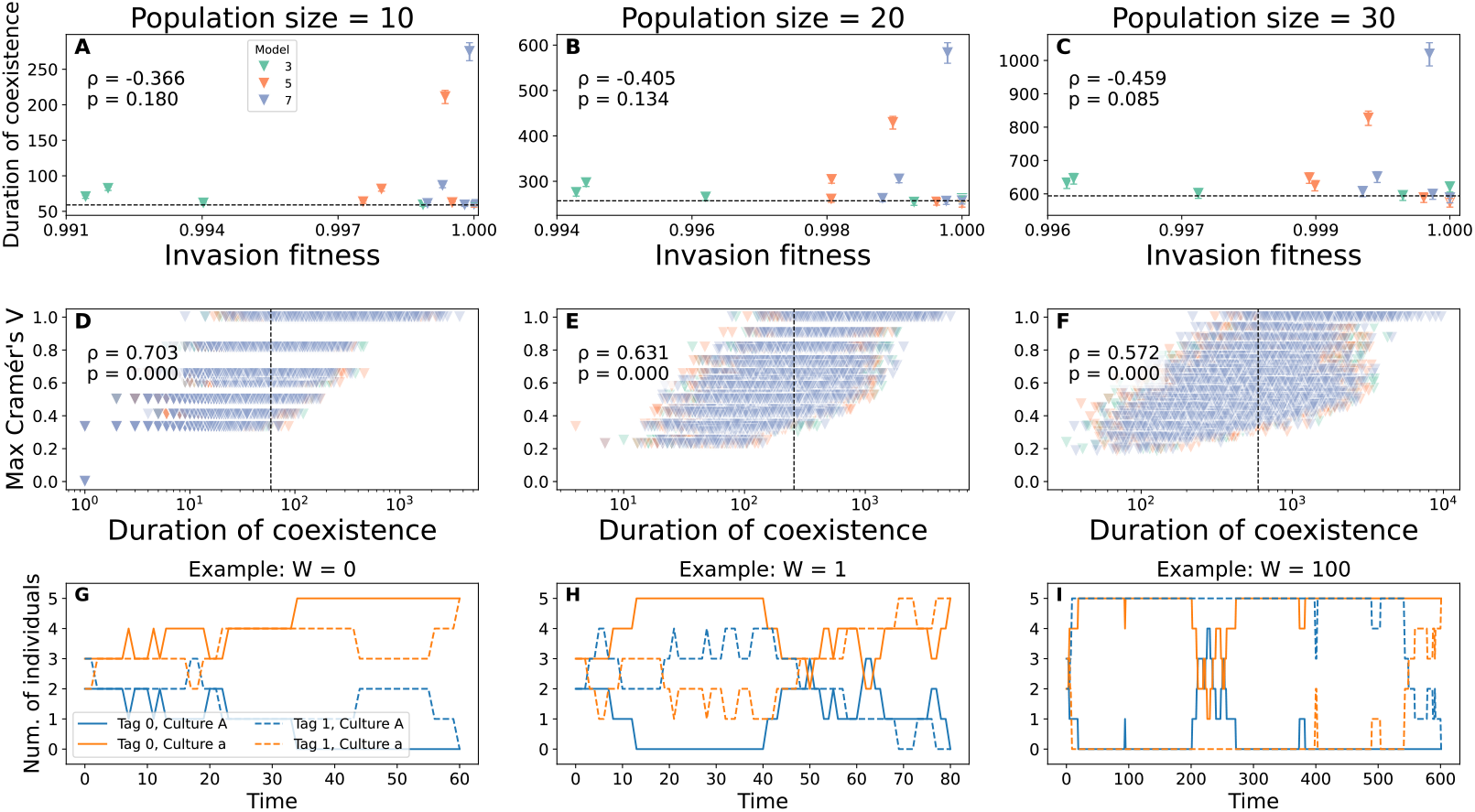
Minimum invasion fitness fails to predict cultural coexistence under the similarity bias Panels A – C: Invasion fitness did not correlate with the mean duration of coexistence under the similarity bias. While the invasion fitness is lower than one, the two cultural traits coexist longer than under the unbiased social learning scenarios (shown by the dashed lines). The colors of the symbols represent the number of cultural models *K*, and the error bars show the standard deviation. Panels D – F: Duration of coexistence was positively correlated with the maximum Cramér’s *V* over time. *V* = 1 means that all individuals with tag 0 own one cultural trait while individuals with tag 1 have the other culture. On panels A –F, each column corresponds to different population size (A and D: *N* = 10, B and E: *N* = 20, and C and F: *N* = 30), and *ρ* and *p* in each panel represent the Spearman’s rank correlation coeffcient and its p-value. Panels G – I: Three typical dynamics when *N* = 10 and *K* = 7 with various strength of the similarity bias *W*. Panel G corresponds to the unbiased social learning *W* = 0, where the mean duration of coexistence is 59. Panel H and I correspond to weak (*W* = 1) and strong (*W* = 100) similarity biases, respectively.

This pattern appeared because the similarity bias generated a substructure in the population: people with tag 0 had one culture, whereas people with tag 1 had another. We measured the degree of the association between the tags and cultural traits based on Cramér’s *V*, where *V* = 0 and *V* = 1 indicate that the cultural traits are independent and perfectly associated with the tags, respectively. Fig. 5D–F shows that the duration of coexistence was longer when the maximum value of Cramér’s *V* over time. This indicated that, once a substructure is generated, the two cultural traits coexist for longer than in unbiased social learning. Figs. S4 and S5 show that the stronger similarity bias tended to increase the duration of coexistence and Cramér’s *V*, respectively. In other words, a stronger similarity bias is more likely to generate a substructure in the population and promote cultural trait coexistence.

Cases of typical dynamics in individual-based simulations have clarified this process. When the similarity bias was absent or weak (Figs. 5G and H), the two cultural traits coexisted until 60–80 time steps, which was close to the mean duration of coexistence under unbiased social learning (59 time steps). Under strong similarity bias (Fig. 5I), the two cultural traits coexisted about 10-times longer (*>* 600 time steps) than under unbiased social learning. In this example, all individuals with tag 0 and 1 had culture *a* and *A*, respectively, at approximately *t* = 30. This condition continued stably until *t* = 550, after which culture *a* spread among individuals with tag 1. In other words, once the cultural traits were associated with the tag traits, neither trait became rare, preventing prediction based on invasion fitness. Consequently, the invasion criterion cannot predict cultural coexistence under similarity bias, either qualitatively or quantitatively.

## 4. Discussion

Cultural diversity is a hallmark of human life. Previous studies have reported the presence of cultural variation within populations (Eberhard et al., 2024; Warf and Vincent, 2007) and that it can drive innovation (Prenzel et al., 2024). However, theoretical studies of the underlying mechanisms are limited. While a previous study analyzed how innovation affects the diversity of competing cultural traits (Kandler and Laland, 2009), we analyzed the duration of multiple cultures coexisting under social learning biases by applying the invasion criterion—a fundamental framework for describing the maintenance of diversity in ecology (Grainger et al., 2019). Our mathematical models showed that the MIF (Schreiber et al., 2023), at least to a moderate degree, predicts cultural coexistence under content-, conformity-, anticonformity-, prestige-, and success-biased social learning, unlike under the similarity bias.

The MIF accurately predicted the mean duration of coexistence of the two cultures under unbiased, content-biased, conformity-biased, and anticonformity-biased social learning (Figs. 2 and S1). While analyzing the inverse of the transition matrix in Eq (13) can derive the mean duration of cultural coexistence (Otto and Day, 2007), our results highlight that only a few elements of the transition matrix (see Eq (1)) are suffcient. This indicates that the invasion criterion is useful when population size *N* is large in simple scenarios. The computational complexity of calculating the inverse of an *N* × *N* matrix is *O*(*N* ^3^) using standard algorithms such as LU decomposition, where *O* (*·*) denotes Landaus Big O notation. However, the invasion fitness requires two elements of the transition matrix for each culture. The MIF also predicted the mean duration of coexistence of three cultural traits (Fig. S2), where deriving the transition matrix was technically diffcult. In addition, the transition matrix derives the mean time until one cultural trait outcompetes the others, which differs from the duration of cultural coexistence when three or more cultural traits coexist. Overall, the invasion criterion is a powerful tool for predicting the mean duration of cultural coexistence in content-biased and frequency-dependent social learning.

Notably, the predictability of MIF was lower in model-biased social learning than in other scenarios. For binary prestige-biased social learning, we calculated the conditional MIF assuming that a low-prestige individual holds a rare cultural trait. Although the mean duration of cultural coexistence increased with conditional MIF, the pseudo-*R*^2^ of the fitted curves was lower and MAPE was higher than for content-biased and frequency-dependent learning (Table 2). This discrepancy may stem from the simplification inherent in using only conditional MIF; high-prestige individuals may also hold rare cultural traits with a non-negligible probability. If we can estimate the probability that individuals with different prestige levels hold a rare trait, we can refine the MIF by weighting the conditional MIF accordingly. However, these probabilities are not analytically tractable and must be obtained through simulations. Another technical challenge arises when prestige is modeled more realistically. Previous studies have analyzed prestige as a continuous or temporally dynamic trait (Egozi and Ram, 2024; Nakata et al., 2024), in which defining an appropriate approximation of the MIF becomes even more diffcult. These complexities highlight the need for refined invasion-based metrics and alternative indicators suitable for prestige-biased social learning.

Under success bias, newborns are more likely to learn from individuals with lower mortality rates, assuming that mortality is negatively correlated with success. In the absence of environmental fluctuations, success bias returns to a special case of context-biased learning, in which a culturally attractive trait lowers the mortality of its holder. In this case, the MIF accurately predicted the mean duration of coexistence between the two cultural traits (Fig. S1). When environmental conditions fluctuated, we assumed that successful cultural traits would change over time. The MIF still predicted the mean duration of coexistence but its accuracy was lower than for content-biased and frequency-dependent learning (Table 2), likely because the MIF was calculated as an average over the environmental states, whereas the stationary distribution of these states does not capture the temporal stability of the environment. The predictive accuracy of MIF decreased as the environmental stability increased, although the average MIF remained unchanged (Fig. S3). Under infrequent environmental changes, a single environmental state persists for a long period, allowing one advantageous culture to exclude the other. Consequently, the actual dynamics deviate from those predicted by the average MIF. In contrast, more frequent environmental changes led to better alignment between the averaged MIF and observed coexistence durations (Table S3). These findings align with the ecological theories of the storage effect, which posit that temporal environmental variations can promote coexistence. A previous study demonstrated that coexistence is favored when environmental changes occur on timescales comparable to key biological rates (Johnson and Hastings, 2022). Therefore, the invasion criterion would be useful under success-biased social learning in the absence of environmental fluctuations or presence of frequent fluctuations.

The MIF failed to predict the duration of coexistence under the similarity-biased social learning scenarios. While the MIF was slightly smaller than one, the duration of cultural coexistence in this scenario was longer than that under unbiased social learning (Figs. 5A–C). Similarity-biased social learning led to an emergent association between cultural and tag characteristics. Once such an association is established, it is unlikely that one cultural trait will spread from people with a focal tag to others with another tag. In other words, one cultural trait is unlikely to be rare when tags and cultural traits are associated (Fig. 5I). Theoretically, with strong similarity bias, our model is similar to the deme-structured population with weak migration, where deme corresponds to populations composed of individuals with identical tag traits among who cultural traits can more easily spread. Hauert et al. (2014) showed that a lower migration rate in a deme-structured population results in longer coexistence. Similarly, our model showed that stronger similarity biases result in longer durations of cultural coexistence (Fig. S4). Overall, our similarity-biased social learning simulations highlight the limitations of the invasion criterion in structured populations (Giaimo et al., 2018).

Nevertheless, the invasion criterion is beneficial for elucidating how social learning biases affect cultural coexistence (Fig. 1). For example, how the strength of social learning biases altered MIF differed between content-, conformity-, and success-biased social learning, although all decreased MIF and the duration of cultural coexistence. On the one hand, the MIF under content- and conformity-biased social learning showed a decreasing curve with a plateau (Figs. 1A and B). On the other hand, the MIF under success-biased social learning showed a nonlinear negative relationship characterized by little change with weak social learning bias, but a sharp decline with stronger bias (Fig. 1E). The effect of increasing the number of cultural models *K* (line colors in Fig. 1) varied among the social learning biases. While it decreased the MIF under content-, conformity-, and prestige-biased social learning biases, this trend was reversed under anticonformity- and success-biased social learning.

These results may be helpful for describing the mechanisms maintaining cultural diversity. Our findings suggest that increasing the number of cultural models, such as by assigning more mentors, may facilitate innovation by maintaining cultural diversity, particularly when individuals exhibit anticonformity- or success-biased social learning. Conversely, content, conformity, and prestige biases may hinder cultural diversity and limit innovation. Beyond the human context, our results may also inform conservation strategies for non-human animals. Socially transmitted behaviors, such as foraging techniques, vocalizations, and migratory routes, are increasingly recognized as part of animal culture, and preserving their cultural diversity can enhance population resilience and adaptive responses to emerging stressors (Brakes et al., 2025). Understanding the social learning biases in animal populations could inform the development of strategies for conserving cultural traits and, by extension, biodiversity.

One limitation of the invasion criterion is that we could not compare the duration of cultural coexistence across social learning biases. Fig.2 shows that content- and conformity-biased social learning resulted in similar MIF, yet their mean duration of cultural coexistence differed. In other words, the identical MIF did not support cultural coexistence for the same duration between the two social learning biases. This echoes previous findings in ecology, where invasion growth rates predicted the coexistence time of multiple species qualitatively but not quantitatively (Pande et al., 2020; Ellner et al., 2020). That is, a larger invasion criterion does not guarantee a longer duration of coexistence. In our case, our model cannot tell whether changing social learning biases can enhance or impede cultural coexistence by simply comparing MIF. Nevertheless, our analysis revealed that MIF can provide qualitative information when social learning biases are fixed in many cases.

In conclusion, this study revealed that the invasion criterion is an appropriate proxy for cultural coexistence under content-, conformity-, anticonformity-, simple prestige-, and success-biased social learning. While evolutionary theory has enriched cultural evolutionary theory, ecological theory can also contribute to its development in terms of how biological diversity is maintained. However, it is not suffcient to simply apply ecological theory to cultural contexts. Therefore, future studies should integrate and extend ecological theory to align it with cultural evolution and develop a unified framework for maintaining diversity.

## Supporting information

Supporing information

## Data availability

Codes and data used in this manuscript are available in the GitHub repository: https://github.com/ShotaSHIBASAKI/CulturalCoexistenceModel.

## Acknowledgment

We would like to thank Editage (www.editage.jp) for English language editing. This study was supported by a grant-in-aid in FY2025 from the Harris Science Research Institute of Doshisha University to S.S., and by NIG-JOINT (60A2025) to S.S. and M.Y.

## Author contribution

S.S.: Conceptualization, Software, Formal analysis, Investigation, Writing - Original Draft, Writing - Review & Editing, Visualization, Funding acquisition. M. Y.: Conceptualization, Writing - Review & Editing, Funding acquisition.

