## Supplementary material for "Applying invasion criterion to cultural evolution": Supporing information

### SI 1 Learning probability is independent of prestige bias strength

In this section, we consider the coexistence of two neutral cultural traits under the binary prestige bias. Consider a population of size $N$ consisting of two prestige classes:

- High prestige individuals: $N_h$
- Low prestige individuals: $N - N_h$.

We denote $w$ as the strength of prestige bias toward the high-prestige individuals, while the bias toward the low-prestige individuals is one. At each time step, we sample $K$ models from the population, among which $M$ are high-prestige individuals, while $K - M$ are low-prestige individuals. Then, a newborn individual inherits a cultural trait from one of $K$ models, depending on the prestige of the models. One individual then dies and is replaced by the newborn individual. The newborn individual inherits the prestige of the dead individual so that the numbers of high and low-prestige people are constant over time.

If we assume each individual can have a rare cultural trait with probability $1/N$ (i.e., the naïve invasion fitness ), the invasion fitness is given as follows:

$$I = \frac{N-1}{N} + L \tag{S15}$$

where $L$ is the probability that a newborn individual learns a rare probability. In this section, we show that $L = 1/N$, i.e., the naïve invasion fitness is identical to that of the unbiased cases in the main text, and that it does *not* depend on the strength of the prestige bias $w$.

The learning probability can be decomposed by two factors, the conditional probability that the focal rare trait is learned by a newborn individual when a high-prestige (low-prestige) individual has the rare cultural trait $L_{\text{high}}$ and that when a low-prestige individual has the rare cultural trait $L_{\text{low}}$:

$$L = \frac{1}{N}\left(N_h L_{\text{high}} + (N - N_h) L_{\text{low}}\right), \tag{S16}$$

where

$$\begin{aligned} L_{\text{high}} &= \sum_{M=1}^{\min\{K,N_h\}} \frac{\binom{N_h-1}{M-1}\binom{N-N_h}{K-M}}{\binom{N}{K}} \frac{(1+w)M}{K+wM} \\ &= \sum_{M=1}^{\min\{K,N_h\}} l_{\text{high}}(M) \frac{1+w}{K+wM}, \end{aligned} \tag{S17}$$

$$\begin{aligned} L_{\text{low}} &= \sum_{M=0}^{\min\{K-1,N_h\}} \frac{\binom{N_h}{M}\binom{N-N_h-1}{K-M-1}}{\binom{N}{K}} \frac{1}{K+wM} \\ &= \sum_{M=0}^{\min\{K-1,N_h\}} l_{\text{low}}(M) \frac{1}{K+wM}. \end{aligned} \tag{S18}$$

Here, $l_{\text{high}}(M)$ and $l_{\text{low}}(M)$ represent the probability that the focal high- or low-prestige individual is chosen as a model by the newborn individual given the number of high-prestige models $M$.

Using Pascal's identity for binomial coefficients, $n\binom{n-1}{k-1} = k\binom{n}{k}$, $N_h L_{\text{high}}$ and $(N - N_h)L_{\text{low}}$ are simplified as follows.

$$N_h L_{\text{high}} = \frac{1}{\binom{N}{K}} \sum_{M=1}^{\min\{K,N_h\}} \binom{N_h}{M}\binom{N-N_h}{K-M} \frac{(1+w)M}{K+wM}, \tag{S19}$$

$$(N - N_h)L_{\text{low}} = \frac{1}{\binom{N}{K}} \sum_{M=0}^{\min\{K-1,N_h\}} \binom{N_h}{M}\binom{N-N_h}{K-M} \frac{K-M}{K+wM} \tag{S20}$$

These two have similar forms but the range of the sum differs. Notice that, however, $l_{\text{high}}(0) = 0$ and that $K - M = 0$ when $M = K$. These two facts allow us to add the two expressions:

$$
\begin{aligned}
N_h L_{\text{high}} + (N - N_h) L_{\text{low}} &= \frac{1}{\binom{N}{K}} \sum_{M=0}^{\min\{K,N_h\}} \binom{N_h}{M} \binom{N-N_h}{K-M} \\
&= \frac{1}{\binom{N}{K}} \binom{N}{K} \\
&= 1.
\end{aligned} \tag{S21}
$$

Hence,

$$
L = \frac{1}{N}. \tag{S22}
$$

**Conclusion:** The learning probability $L$ does not depend on the prestige bias strength $w$ and is always equal to $\frac{1}{N}$. This indicates that the naïve invasion growth rate in the binary prestige model is always identical to the unbiased case in the main text.

### SI 2 When cultural traits affect the mortality rates

In this supporting information, we analyzed the scenarios where having cultural trait $A$ increased or decreased the morality rate of owners. While the morality of owners of cultural trait $a$ was fixed as 1, we varied the mortality rate of owners of cultural trait $A$ as $1 + d$ where $d = \{\pm 0.2, \pm 0.1, \pm 0.01, 0\}$. The results of unbiased, content-biased, (anti)conformity-biased, prestige-biased, and similarity-biased social learning in the main text corresponded to $d = 0$. Under the success-biased social learning, we incorporated the variation in $d$ to represent that successful individuals had smaller mortality.

In the presence of the cultural impacts on the mortality rate, one of $n_A$ individuals with cultural trait $A$ is dead at each time step with the probability of $n_A d/(N + n_a d)$. Then, the invasion fitness of cultural trait $A$ is rewritten as follows:

$$
I_A = 1 + \frac{(N-1)Kl_A(n_A = 1; K)}{(N+d)N} - \frac{1+d}{N+d}\left(1 - \frac{Kl_A(n_A = 1; K)}{N}\right). \tag{S23}
$$

In this section, we analyzed whether the cultural impact on the mortality rate affects the prediction by the modern coexistence theory in scenarios of the unbiased, content-biased, and (anti)conformity social learning biases. These scenarios were easily analyzed using the Markov chain model. The transition matrix $P$ is written as follows:

$$
P_{ij} \equiv P(N_A(t+1) = j | N_A(t) = i) = \begin{cases} \frac{(N-i)p_A(i)}{N+id} & \text{if } j = i+1 \\ \frac{i(1+d)(1-p_A(i))}{N+id} & \text{if } j = i-1 \\ \frac{N(1-p_A(i)) + i(2p_A(i) - 1 + dp_A(i))}{N+id} & \text{if } j = i \\ 0 & \text{otherwise.} \end{cases} \tag{S24}
$$

Fig. S1 shows that the invasion fitness positively correlates with the mean duration of coexistence under the unbiased, content-biased, and (anti) conformity-biased social learning when the cultural trait affects the mortality.

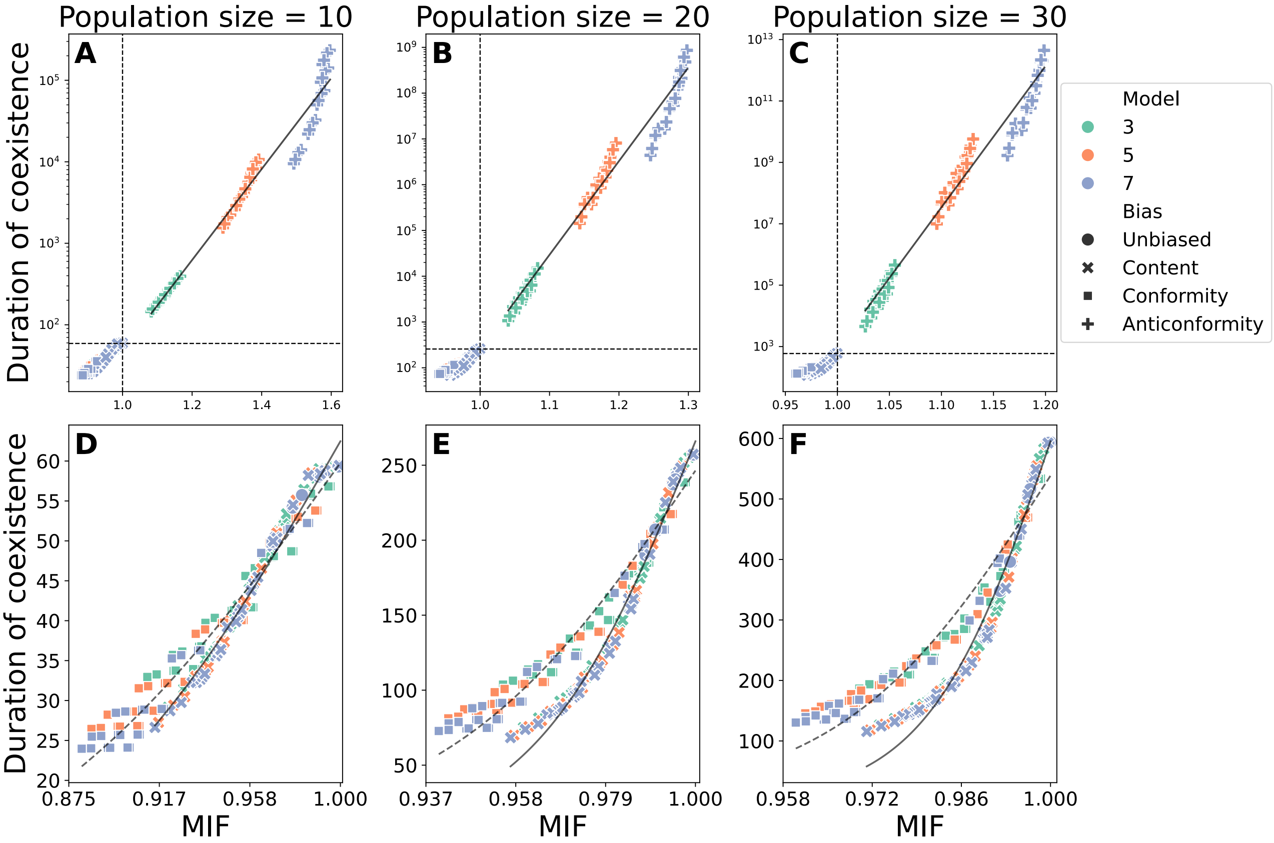


Figure S1: MIF predicts the coexistence of non-neutral cultural traits under unbiased, content-biased, and frequency-dependent biased social learning

Similar to Fig. 2, each column corresponds to a different population size $N$. Panels D – F show the lower-left part in panels A – C, respectively. The current figure allows cultural trait $A$ to increase or decrease the mortality of its owner. The statistical details are shown in Table S1.

Table S1: Summary of regression analysis when cultural traits affect mortality rates

| Bias | $N$ | Model | (pseudo-)$R^2$ | MAPE |
|---|---|---|---|---|
| Content | 10 | $y = \frac{124.95}{1+\exp\{-15.21(x-1)\}}$ | 0.98 | 2.16 |
| | 20 | $y = \frac{531.93}{1+\exp\{-52.73.(x-1)\}}$ | 0.98 | 5.23 |
| | 30 | $y = \frac{1190.70}{1+\exp\{-102.37(x-1)\}}$ | 0.96 | 11.21 |
| Conformity | 10 | $y = \frac{119.53}{1+\exp\{-12.66(x-1)\}}$ | 0.96 | 5.16 |
| | 20 | $y = \frac{492.82}{1+\exp\{-33.74(x-1)\}}$ | 0.96 | 6.99 |
| | 30 | $y = \frac{1077.02}{1+\exp\{60.25(x-1)\}}$ | 0.95 | 10.11 |
| Anticonformity | 10 | $\log_{10} y = 5.58x - 3.90$ | 0.97 | 2.63 |
| | 20 | $\log_{10} y = 20.46x - 18.04$ | 0.98 | 3.28 |
| | 30 | $\log_{10} y = 46.02x - 43.09$ | 0.98 | 4.21 |

$x$: MIF, $y$: mean duration of coexistence.

### SI 3 Models with three-cultural traits

In this section, we analyzed the mean duration of coexistence of three cultural traits under the unbiased, content-biased, conformity-biased, and anticonformity-biased social learning. We applied the individual-based model with $N = 30$ and $K = 3, 5, 7$, because the analysis of the Markov chain model is technically hard under the three or more cultural traits scenarios. We modified the strength of the social learning biases as in the main text. Under the content-biased social learning, we assumed that the preference for one culture was fixed as one, the second was given by $1 + s$, and the third one was $1 + 2s$. We also allowed the cultural traits to change the mortality of their owners. While one culture did not affect the mortality, the second culture additively changed the mortality of its owner by $d$, and the third culture changed by $2d$. The value of $d$ was identical to SI 2. At the initial state, each individual has either of three three cultural traits with probability of 1/3.

To calculate the invasion fitness, we ran simulations of the rest two cultural scenarios until $t = 90$ as in the main text, after which we randomly chose one individual from the population and gave him/her the new focal cultural trait. This led the realized invasion fitness, given the number of owners of the two residential traits. By repeating the above calculation 1,000 times, we defined the invasion fitness of the focal culture as the mean of the 1,000 realized invasion fitness. Fig. S2 and Table S2 show that the MIF accurately predicts the mean duration of three cultural traits.

The invasion fitness of a focal culture depends on the number of the owners of the two resident cultures.

Table S2: Summary of regression analysis in three-culture models

| Bias | $N$ | Model | (pseudo-)$R^2$ | MAPE |
|---|---|---|---|---|
| Content | 30 | $y = \frac{480.56}{1+\exp\{-36.6(x-1)\}}$ | 0.64 | 13.66 |
| Conformity | 30 | $y = \frac{441.73}{1+\exp\{-36.48(x-1)\}}$ | 0.94 | 6.41 |
| Anticonformity | 30 | $\log_{10} y = 17.32x - 15.29$ | 0.95 | 4.52 |

$x$: MIF, $y$: mean duration of coexistence.

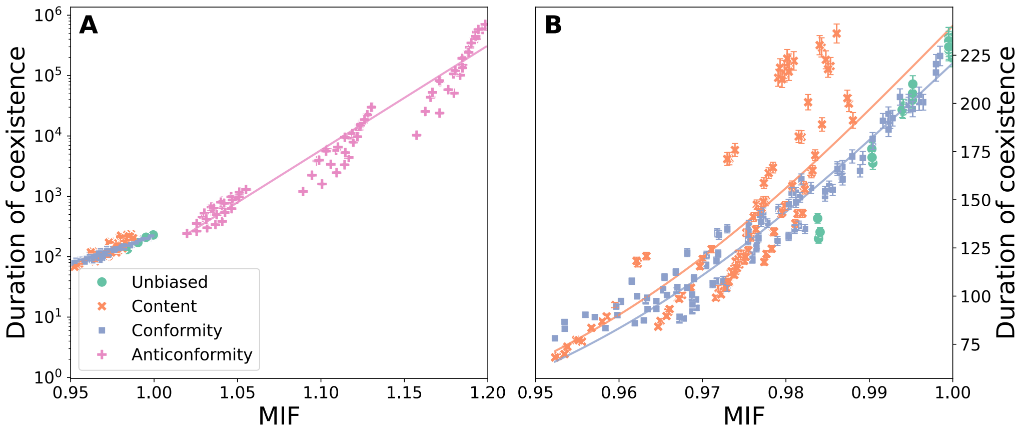


Figure S2: MIF predicts the coexistence of three cultural traits

The MIF predicts the coexistence of three cultural traits under the unbiased (green circles), content-biased (orange x marks), conformity-biased (blue squares), and anticonformity-biased (pink cross marks) social learning. Each symbol corresponds to the mean duration of coexistence from 1,000 replicates, and the error bars show the standard errors. Panel B corresponds to the lower left part of panel A. The colored lines represent the statistical models fitted to the simulation data from each social learning bias. Table S2 provides the statistical details.

### SI 4 Supporting figures and tables

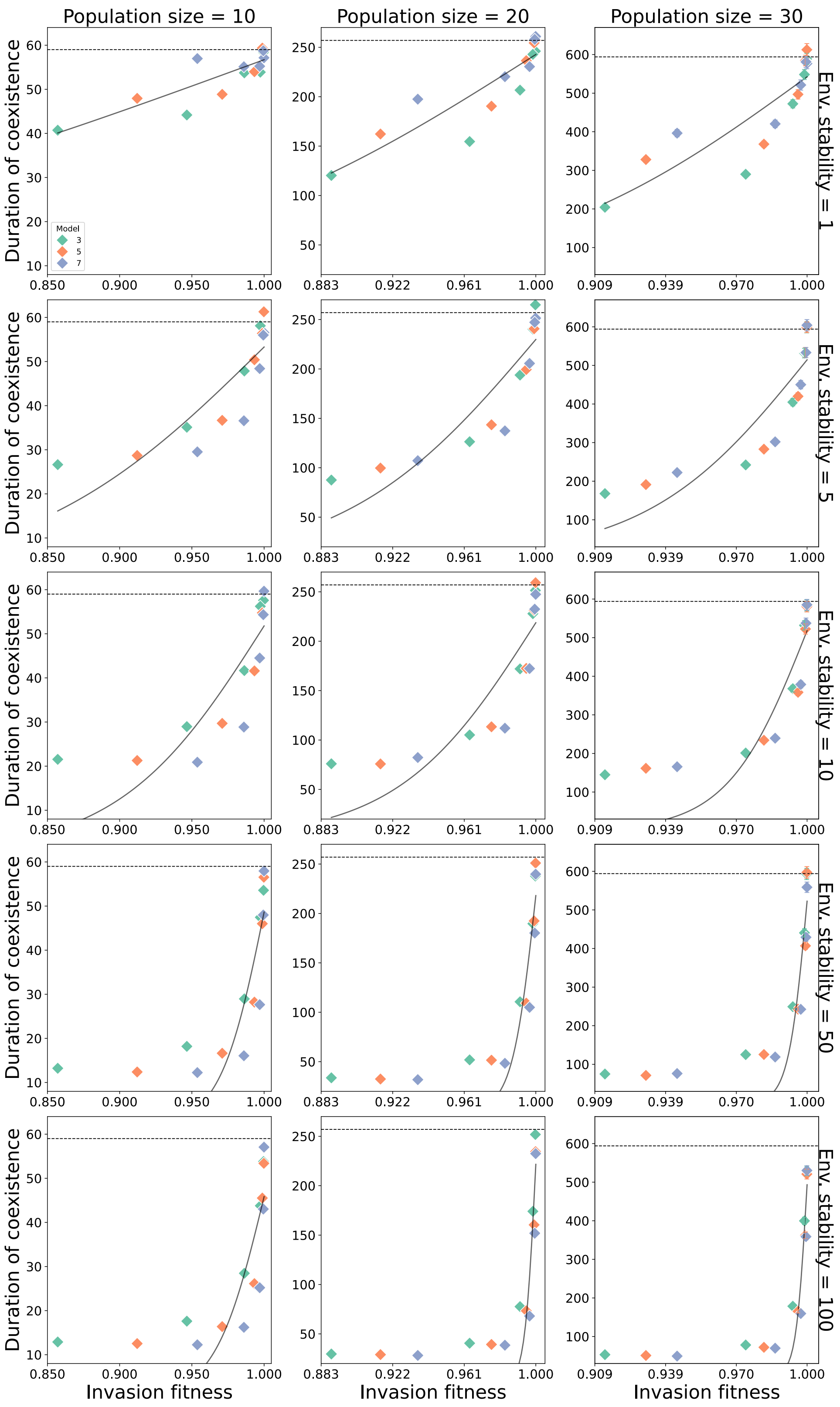


Figure S3: Success bias scenario at each environmental stability

The invasion fitness and the mean duration of the coexistence of the two cultural traits under the success bias with environmental fluctuations. Each row corresponds to the environmental stability, while each column corresponds to different population size. The horizontal dashed lines represent the mean duration of coexistence without social learning biases (Fig. 2).

Table S3: Summary of regression analysis under success bias

| $N$ | Env. stability | coeff. $a$ | coeff. $b$ | pseudo-$R^2$ | MAPE |
|---|---|---|---|---|---|
| 10 | 1 | 113.41 | 4.22 | 0.72 | 4.34 |
| 20 | 1 | 485.48 | 9.75 | 0.79 | 7.42 |
| 30 | 1 | 1084.63 | 16.14 | 0.74 | 12.77 |
| 10 | 5 | 106.597 | 12.05 | 0.73 | 12.76 |
| 20 | 5 | 459.68 | 19.07 | 0.80 | 15.50 |
| 30 | 5 | 1028.97 | 28.92 | 0.77 | 21.19 |
| 10 | 10 | 103.57 | 65.90 | 0.69 | 21.77 |
| 20 | 10 | 437.39 | 26.61 | 0.76 | 28.18 |
| 30 | 10 | 1036.18 | 58.51 | 0.75 | 28.97 |
| 10 | 50 | 97.24 | 65.90 | 0.69 | 39.28 |
| 20 | 50 | 436.16 | 155.50 | 0.84 | 44.36 |
| 30 | 501045.18 | | 248.94 | 0.83 | 48.00 |
| 10 | 100 | 91.68 | 58.68 | 0.66 | 38.88 |
| 20 | 100 | 432.32 | 334.39 | 0.83 | 54.36 |
| 30 | 100 | 986.67 | 421.25 | 0.88 | 52.00 |

$x$: MIF, $y$: mean duration of coexistence.

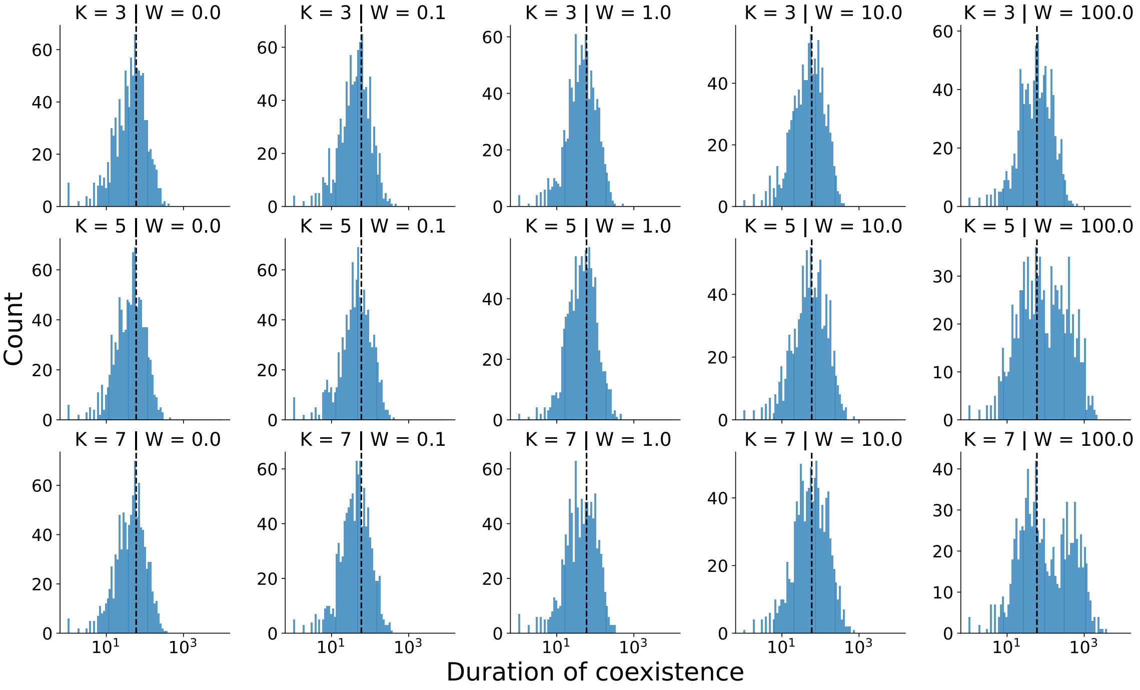


Figure S4: Strength of similarity bias increases the duration of coexistence

Distributions of coexistence under the similarity bias at each number of cultural models $K$ and the strength of the similarity bias $W$ are shown, while the population size was fixed at $N = 10$. In each panel, we ran the agent-based model 1000 times. The dashed lines represent the mean duration of coexistence under the unbiased social learning ($W = 0$).

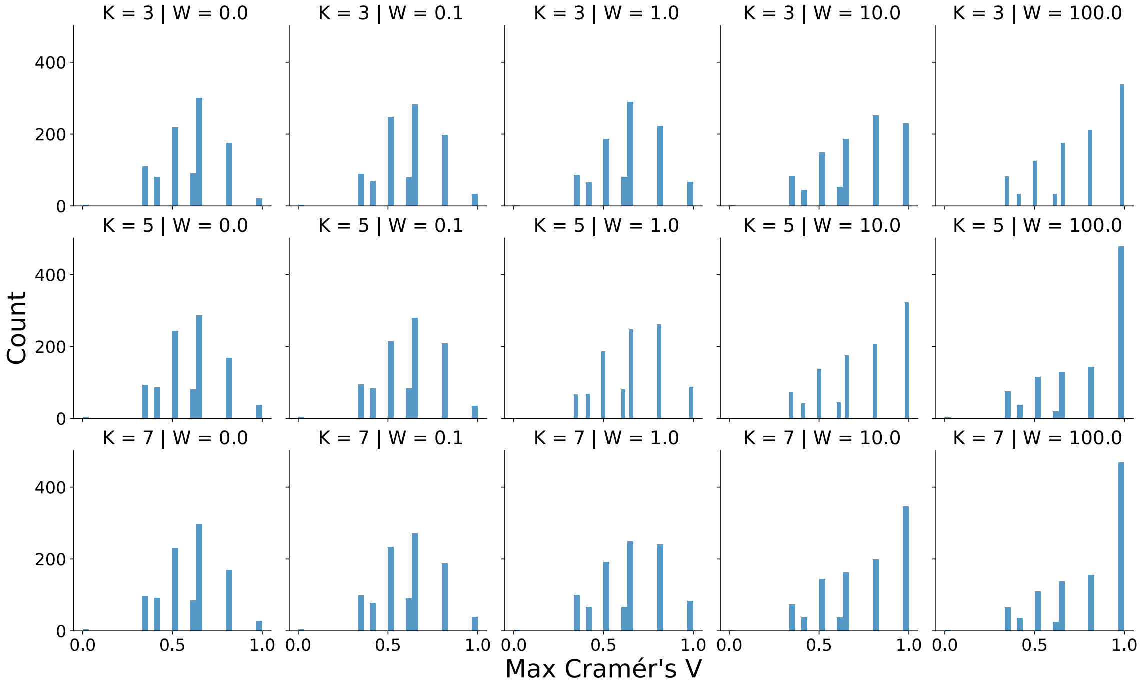


Figure S5: Strength of similarity bias increases the Cramér's V

Distributions of maximum Cramér's V in each run at each number of cultural models $K$ and the strength of the similarity bias $W$ are shown, while the population size was fixed at $N = 10$. In each panel, we ran the agent-based model 1000 times.
